# Genetic Contributions To Health Literacy

**DOI:** 10.1101/595967

**Authors:** Chloe Fawns-Ritchie, Gail Davies, Saskia P Hagenaars, Ian J Deary

**Affiliations:** Centre for Cognitive Ageing and Cognitive Epidemiology, University of Edinburgh, Edinburgh, UK, EH8 9JZ; Department of Psychology, University of Edinburgh, Edinburgh, UK, EH8 9JZ; Social, Genetic and Developmental Psychiatry Centre, Institute of Psychiatry, King’s College London, London, UK, SE5 8AF

**Author notes:** Corresponding author: Ian J Deary, Department of Psychology, University of Edinburgh, 7 George Square, Edinburgh, EH8 9JZ, Scotland, UK.

## Abstract

Higher health literacy is associated with higher cognitive function and better health. Despite its wide use in medical research, no study has investigated the genetic contributions to health literacy. Using 5,783 English Longitudinal Study of Ageing (ELSA) participants (mean age=65.49, SD=9.55) who had genotyping data and had completed a health literacy test at wave 2 (2004-2005), we carried out a genome-wide association study (GWAS) of health literacy. We estimated the proportion of variance in health literacy explained by all common single nucleotide polymorphisms (SNPs). Polygenic profile scores were calculated using summary statistics from GWAS of 21 cognitive and health measures. Logistic regression was used to test whether polygenic scores for cognitive and health-related traits were associated with having adequate, compared to limited, health literacy. No SNPs achieved genome-wide significance for association with health literacy. The proportion of variance in health literacy accounted for by common SNPs was 8.5% (SE=7.2%). Greater odds of having adequate health literacy were associated with a 1SD higher polygenic score for general cognitive ability (OR=1.34, 95% CI 1.26-1.42), verbal-numerical reasoning (OR=1.30, 1.23-1.39), and years of schooling (OR=1.29, 1.21-1.36). Reduced odds of having adequate health literacy were associated with higher polygenic profiles for poorer self-rated health (OR=0.92, 0.87-0.98) and schizophrenia (OR=0.91, 0.85-0.96). The well-documented associations between health literacy, cognitive function and health may partly be due to shared genetic aetiology. Larger studies are required to obtain accurate estimates of SNP-based heritability, and to discover specific health literacy-associated genetic variants.

## Introduction

Health literacy is “the degree to which individuals have the capacity to obtain, process, and understand basic health information and services needed to make appropriate health decisions”.^1^ This capacity is thought to be important for navigating all aspects of health care, including the ability seek out and act upon appropriate health information, and self-manage health conditions.^1, 2^ Tests of functional health literacy have been used to investigate the association between health literacy and health. Individuals with lower health literacy have been found to be less likely to take part in health promoting behaviours.^3^ Lower health literacy is associated with poorer overall health status,^4^ lower self-reported physical and mental health,^3, 5^ and greater self-reported depressive symptoms.^6^ One study^5^ found that individuals with inadequate health literacy were 48% more likely to report a diagnosis of diabetes and 69% more likely to report having heart failure, compared to those with adequate health literacy, after adjusting for sociodemographic variables and health behaviours. Using prospective studies, lower health literacy predicted incident dementia^7, 8^ and risk of dying.^4, 9, 10^

Compared with those of health literacy, similar associations with health have been found for cognitive function. Individuals with higher cognitive function tend to participant more in health promoting behaviours.^11–14^ Vascular risk factors, including diabetes and hypertension, have been associated with poorer cognitive function and greater cognitive decline.^15–18^ Cognitive function, measured early in life, has been found to predict later life physical functioning and health status,^19^ psychological distress,^20^ psychiatric illness,^21–25^ dementia,^26^ and death.^27–30^

Performance on tests of health literacy and cognitive function are moderately to highly correlated.^31–34^ Murray et al.^32^ found that the correlations between general cognitive ability and three tests of health literacy, tested in older adulthood, ranged from 0.35 to 0.53 (p <.001). Given these correlations, researchers have sought to determine whether the relationship between health literacy and health remained when also accounting for cognitive function. Cognitive function has been consistently found to attenuate the size of the association between health literacy and health; however, whereas some studies have found that health literacy no longer predicted better health when controlling for cognitive function,^33, 35–37^ others have found that a small but significant association remained between higher health literacy scores and better health when also controlling for cognitive function.^7, 10, 33, 38, 39^

Whereas there is a wealth of evidence reporting a relationship between health literacy, cognitive function and health, it is less well understood why these associations are found. One possibility is that they share genetic influences. Cognitive function is substantially heritable.^40–42^ With increasing samples sizes, the specific genetic variants associated with cognitive function are being identified.^43, 44^ One study^45^ sought to explore the shared genetic architecture between cognitive function and health, using two complementary genetic techniques: linkage disequilibrium (LD) score regression^46^ and polygenic profile scoring.^47^ The first technique involves calculating the genetic correlations between two traits of interest using summary results from previous GWAS. The second technique uses summary GWAS data for a specific trait (e.g., type 2 diabetes) and tests whether the genetic variants found to be associated with this trait are also associated with the same (e.g., type 2 diabetes) or a different (e.g., cognitive function) phenotype in an independent sample. Using these techniques, Hagenaars et al.^45^ found substantial shared genetic influences between cognitive function and physical and mental health. Negative genetic correlations were found between a test of verbal-numerical reasoning and Alzheimer’s disease (*r_g_* = −0.39, p = 0.002), and schizophrenia (*r_g_* = −0.30, p = 3.5×10^-11^), among others. Polygenic profiles for various mental and physical health-related variables were associated with performance on tests of cognitive function, including coronary artery disease, Alzheimer’s disease, and schizophrenia. The shared genetic architecture between cognitive function and health has been subsequently replicated using larger samples.^44^

Summaries are available regarding the advances made in understanding the genetic architecture of cognitive function and its overlap with physical and mental health.^42, 48^ However, to the best of our knowledge, no one has investigated the genetic contributions to people’s differences in health literacy. The aim of the present study was to explore the genetic contributions to health literacy and its overlap with cognitive function and health. Using data from ELSA, a sample of English adults aged 50 years and older, the present study: conducted a GWAS of health literacy; estimated its SNP-based heritability; and used polygenic profile scoring to examine the genetic overlap between health literacy and cognitive function, and health literacy and various health-related traits.

## Materials and methods

### Participants

This study used data from ELSA (https://www.elsa-project.ac.uk/), a cohort study designed to be representative of English adults aged 50 years and older.^49^ The original sample (wave 1) was recruited in 2002-2003 and consisted of 11,391 participants. Participants have been followed-up every two years and the sample has been refreshed at subsequent waves to ensure the sample’s representativeness. Interviews took place, via computer assisted personal interviews and self-completion questionnaires, in the participants’ own homes. The topics assessed included health, financial and social circumstances. A nurse visit was carried out every second wave to measure biomarkers. Blood samples collected during the nurse visit have been used to genotype ELSA participants. More information on the ELSA sampling procedures are reported elsewhere.^49^ For the present study, a subsample of participants were used who completed the health literacy test at wave 2 (2004-2005) and who had genome-wide genotyping data (n = 5,783).

### Procedure

#### Health literacy

Health literacy was measured using a four-item reading and comprehension test. This test was designed to mimic written materials, such as drug labels, that would be encountered in a health-care setting. A piece of paper containing instructions for an over-the-counter packet of medicine was given to participants. Participants were asked four questions about the information on the medicine packet (e.g., “what is the maximum number of days you may take this medication?”). One point was awarded for each correct answer. As has been done in other studies,^50, 51^ participants were categorised as having adequate (4/4 questions correct) or limited health literacy (<4 correct).

#### Genotyping and quality control

A total of 7,597 ELSA participants who had provided blood samples were genotyped, in two batches (batch one n = 5,652; batch two n = 1,945) by UCL Genomics using the Illumina Omni 2.5-8 chip. Quality control procedures were performed by UCL Genomics and by the present authors. This included removal of SNPs based on call rate, minor allele frequency and deviation from Hardy-Weinberg equilibrium. Individuals were removed based on call rate, relatedness, gender mismatch, and non-Caucasian ancestry. A sample of 7,358 participants remained following quality control procedures.

#### Imputation

Pre-phasing and imputation to the 1000 Genome Phase 3 reference panel^52^ was performed using the Sanger Imputation Service,^53^ EAGLE2 (v2.0.5),^54^ and PBWT^55^ pipeline.

#### Curation of summary results from GWAS of cognitive and health-related traits

Summary results from 21 GWAS of cognitive function, general health status variables, chronic diseases, health behaviours, neuro-psychiatric disorders, years of schooling, social deprivation and the personality traits of conscientiousness and neuroticism were collected. For each trait, we checked the samples used in the GWAS to ensure ELSA was not included. Sources of summary statistics, and key references are given in the Supplementary materials and Supplementary Table 1.

### Statistical analyses

#### Genome-wide association analyses

SNP-based association analyses were performed using the BGENIE v1.2 analysis package (https://jmarchini.org/bgenie/). A linear SNP association model was tested which accounted for genotype uncertainty. Prior to these analyses the health literacy phenotype was adjusted for the following covariates: age, sex, and 15 principal components.

#### Genomic risk loci characterisation using FUMA

Genomic risk loci were defined from the SNP-based association results, using FUnctional Mapping and Annotation of genetic associations (FUMA).^56^ Firstly, independent significant SNPs were identified using the SNP2GENE function and defined as SNPs with a p-value of < 1×10^-5^ and independent of other genome-wide suggestive SNPs at r2 < 0.6. Using these independent significant SNPs, tagged SNPs to be used in subsequent annotations were identified as all SNPs that had a MAF ≥ 0.0005 and were in LD of r2 ≥ 0.6 with at least one of the independent significant SNPs. These tagged SNPs included those from the 1000 Genomes Phase 3 reference panel and need not have been included in the GWAS performed in the current study. Genomic risk loci that were 250 □ kb or closer were merged into a single locus. Lead SNPs were also identified using the independent significant SNPs and were defined as those that were independent from each other at r2 < 0.1.

#### Comparison to previous findings

A look-up of the independent significant and tagged SNPs for health literacy in the current study was performed in previous GWAS of general cognitive ability^44^ and years of schooling.^57^ We identified whether significant SNPs and tagged SNPs reported here reached either genome-wide (p < 5×10^-8^) or suggestive (p < 1×10^-5^) significance in these previous GWAS.

#### Gene-based analysis implemented in FUMA

Gene-based association analyses were conducted using MAGMA.^58^ The test carried out using MAGMA, as implemented in FUMA, was the default SNP-wise test using the mean *χ*^2^ statistic derived on a per gene basis. SNPs were mapped to genes based on genomic location. All SNPs that were located within the gene-body were used to derive a p-value describing the association found with health literacy. The SNP-wise model from MAGMA was used and the NCBI build 37 was used to determine the location and boundaries of 18,199 autosomal genes. LD within and between each gene was gauged using the 1000 genomes phase 3 release. A Bonferroni correction was applied to control for multiple testing across 18,199 genes; the genome-wide significance threshold was p□<□2.75×10^-6^.

#### Functional annotation implemented in FUMA

The independent significant SNPs and those in LD with the independent significant SNPs were annotated for functional consequences on gene functions using ANNOVAR^59^ and the Ensembl genes build 85. Functionally-annotated SNPs were then mapped to genes based on physical position on the genome, and chromatin interaction mapping (all tissues). Intergenic SNPs were mapped to the two closest up- and down-stream genes which can result in their being assigned to multiple genes.

#### Gene-set analysis implemented in FUMA

In order to test whether the polygenic signal measured in the GWAS clustered in specific biological pathways, a competitive gene-set analysis was performed. Gene-set analysis was conducted in MAGMA^58^ using competitive testing, which examines if genes within the gene set are more strongly associated with health literacy than other genes. A total of 10,675 gene-sets (sourced from Gene Ontology,^60^ Reactome,^61^ and, SigDB^62^) were examined for enrichment of health literacy. A Bonferroni correction (p < 0.05/10,675 = 4.68×10^-6^) was applied to control for the multiple tests performed.

#### Gene-property analysis implemented in FUMA

A gene-property analysis was conducted using MAGMA in order to indicate the role of particular tissue types that influence differences in health literacy. The goal of this analysis was to test if, in 30 broad tissue types and 53 specific tissues, tissue-specific differential expression levels were predictive of the association of a gene with health literacy. Tissue types were taken from the GTEx v6 RNA-seq database^63^ with expression values being log2 transformed with a pseudocount of 1 after winsorising at 50, with the average expression value being taken from each tissue. Multiple testing was controlled for using a Bonferroni correction (p < 0.05/53 = 9.43×10^-4^).

#### Estimation of SNP-based hentability

The proportion of variance explained by all common SNPs was estimated using univariate genome-wide complex trait analysis (GCTA-GREML).^64^ The sample size for the GCTA-GREML is slightly smaller than that used in the association analysis (n = 5,661), because one individual was excluded from any pair of individuals who had an estimated coefficient of relatedness of > 0.025 to ensure that effects due to shared environment were not included. The same covariates were included in the GCTA-GREML as for the SNP-based association analysis.

#### Polygenic profile analyses

Polygenic profile scores were created using PRSice version 2^65^ (https://github.com/choishingwan/PRSice). First, we used the GWAS results for health literacy to create health literacy polygenic profile scores in an independent sample, and used these scores to predict health literacy, cognitive function and educational attainment phenotypes. Polygenic profile scores for health literacy were created in 1,005 genotyped participants from the Lothian Birth Cohort 1936 (LBC1936) study^66^ by calculating the sum of alleles associated with health literacy across many genetic loci, weighted by the effect size for each loci. Before the polygenic scores were created, SNPs with a minor allele frequency of < 0.01 were removed and clumping was used to obtain SNPs in LD (*r*^2^ < 0.25 within a 250 kb window). Five scores were then created that included SNPs according to the significance of the association with health literacy, based on the following p-value thresholds: p < 0.01, p < 0.05, p < 0.1, p < 0.5, and all SNPs. Linear regression was used to investigate whether polygenic profiles for health literacy were associated with performance on the Newest Vital Sign,^67^ a test of health literacy similar in content to the ELSA health literacy test, a measure of general cognitive ability, and years of schooling (see Supplementary Methods for more detail on these phenotypes). Models were adjusted for age, sex, and 4 genetic principal components and standardised betas were calculated.

Next, we used summary GWAS results from 21 GWAS of cognitive and health-related phenotypes to create polygenic profile scores for cognitive and health-related traits in ELSA participants. As the creation of polygenic scores requires summary GWAS results from an independent sample, the GWAS of general cognitive ability^44^ was re-run removing ELSA participants. SNPs with a minor allele frequency of < 0.01 were removed and clumping was used to obtain SNPs in LD (*r*^2^ < 0.25 within a 250 kb window) prior to the creation of the polygenic scores. Five scores were created for each phenotype based on the p-value thresholds detailed above. For Alzheimer’s disease, we created a second set of scores with a 500 kb region around the *APOE* locus removed (hereafter called ‘Alzheimer’s disease (500 kb)’) to create a polygenic risk score of Alzheimer’s disease with and without the *APOE* locus.

These polygenic scores were converted to z-scores. Logistic regression was used to investigate whether polygenic profiles for cognitive and health-related traits were associated with having adequate, compared to limited, health literacy in ELSA participants. All models were adjusted for age at wave 2, sex, and the 15 genetic principal components to control for population stratification. For each phenotype, five logistic regression models were run using the five polygenic scores created based on the p-value thresholds; thus, a total of (5×21) 105 models were run. To control for multiple testing, the reported p-values are false discovery rate-corrected. This method controls for the number of false positive results in those that reach significance.^68^ A multivariate logistic regression model was run including all of the significant polygenic scores, controlling for age, sex, and 15 genetic principal components, to test whether these polygenic scores independently contributed to health literacy.

## Results

Of the 7,358 participants who remained following genotyping quality control procedures, 5,783 (3,160 female; 54.6%) had completed the health literacy test at wave 2 and form the analytic sample (mean age = 65.49, SD = 9.55). A total of 4,012 (69.4%) participants had adequate health literacy, whereas 1,771 (30.6%) participants had limited health literacy. Participants with limited health literacy were older (mean age = 67.76, SD = 10.00) than participants with adequate health literacy (mean age = 64.72, SD = 9.19; t (3140.90) = 10.91, p < .001).

### Genome-wide association study

A genome-wide association analysis of health literacy found no genome-wide significant (p < 5×10^-8^) SNP associations. There were 131 suggestive SNP associations (p < 1×10^-5^). The SNP-based Manhattan plot is shown in Figure 1 (the SNP-based QQ plot is shown in Supplementary Figure 1; Suggestive SNPs are reported in Supplementary Data 1). Genomic risk loci characterization performed using FUMA with the genome-wide suggestive significance threshold (p < 1×10^-5^) identified 39 ‘independent’ significant SNPs distributed within 36 loci; see Methods section for description of independent SNP selection criteria. For consistency, we use the term ‘independent suggestively significant SNP’ here according to the definition that is used in the relevant analysis package and the significance threshold described above. Details of functional annotation of these independent suggestively significant SNPs and tagged SNPs within the 36 loci can be found in Supplementary Data 2.

**Figure 1.**
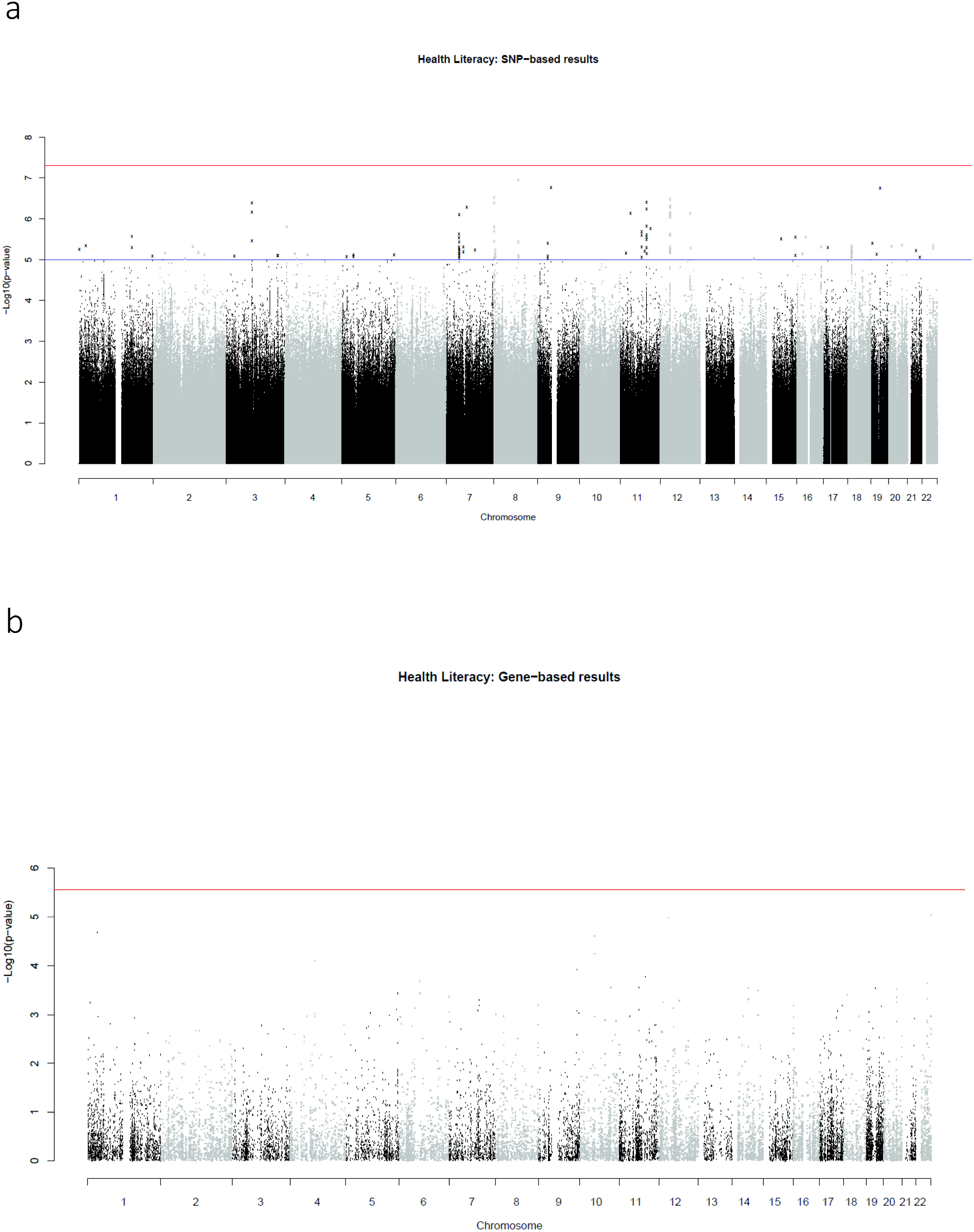
SNP-based (a) and gene-based (b) association results for health literacy. The red line indicates the threshold for genome-wide significance: p<5×10^-8^ for (a), p<2.75×10^-6^ for (b); the blue line in (a) indicates the threshold for suggestive significance: p<1×10^-5^

### Comparison with previous findings

Of the 39 independent suggestively significant and 253 tagged SNPs (those in LD with the independent suggestively significant SNPs), none had been reported as reaching genome-wide (p < 5×10^-8^) or suggestive (p < 1×10^-5^) significance in previous GWAS of general cognitive ability or years of education.

### Gene-based analyses

No genome-wide significant findings were found from the gene-based association analysis; the gene-based association results are shown in Supplementary Data 3 (the gene-based Manhattan plot is shown in Figure 1; the QQ plot is shown in Supplementary Figure 1). The gene-set and gene-property analyses also did not identify any significant results (Supplementary Data 4 and 5).

### SNP basedheritability

We estimated the proportion of variance explained by all common SNPs to be 0.085 (SE = 0.072). We note that, with the large standard error, this does not rule out zero SNP-based heritability.

We did not calculate genetic correlations between health literacy and those phenotypes included in the polygenic profile analyses as we did not have adequate power in this sample to utilise either the LD score regression method or, for those phenotypes also available in ELSA, bivariate GCTA-GREML. The mean chi^2^ value for the health literacy phenotype was 1.009 which is below the LD score regression recommended threshold of 1.02.^46^ This indicates that there is too small a polygenic signal for these methods to work with.

### Health literacy polygenic profile scores predicting health literacy, cognitive function, and educational attainment in LBC1936

Polygenic profile score for health literacy did not significantly predict performance on the Newest Vital Sign, cognitive ability, or years of schooling in LBC1936 (Supplementary Table 2).

### Cognitive and health-related polygenic scores predicting health literacy in ELSA

Table 1 shows the results for the association between cognitive and health-related polygenic scores and health literacy in ELSA participants, using the most predictive threshold. Supplementary Table 3 reports the full results for all thresholds.

**Table 1.**
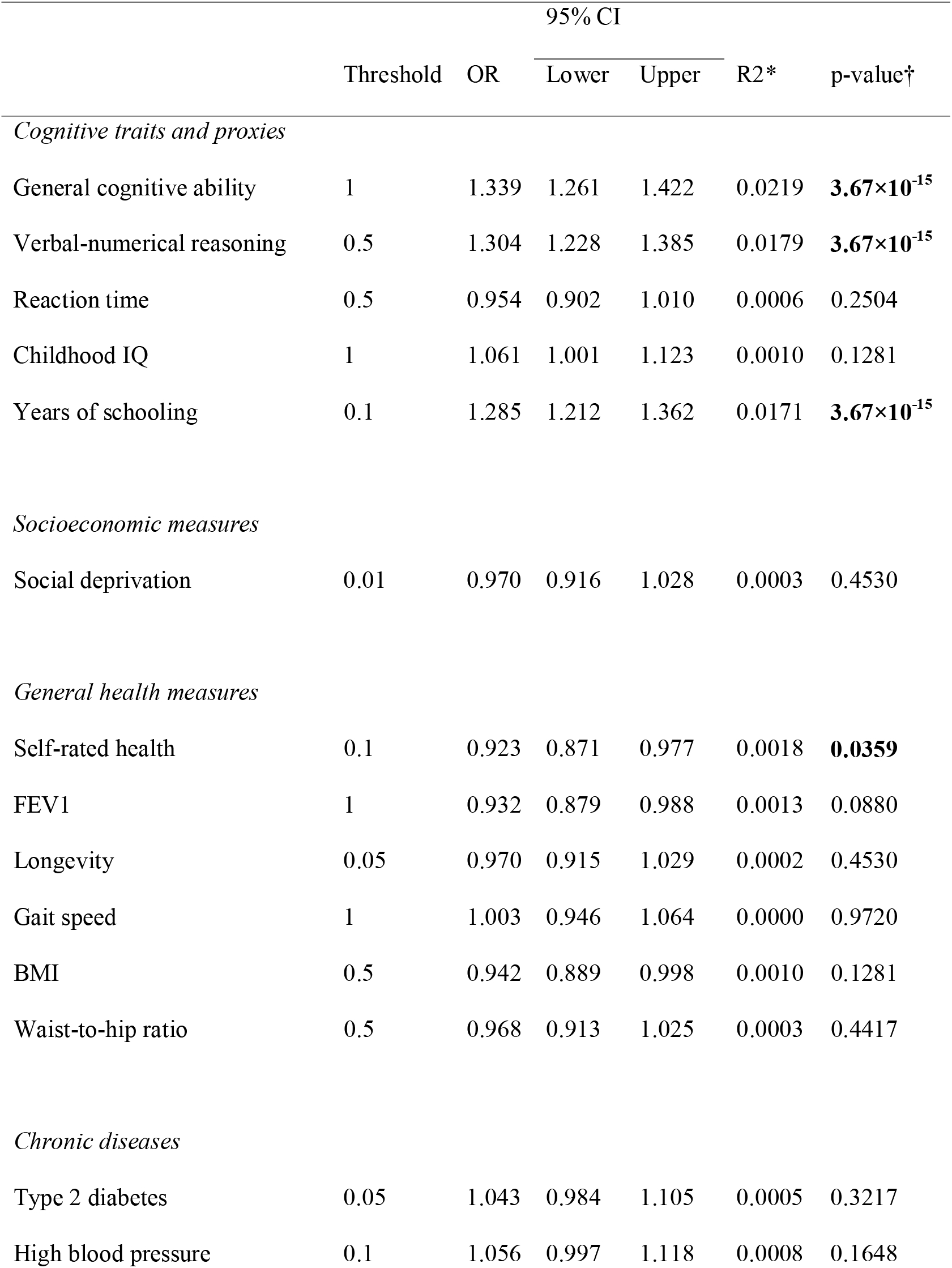
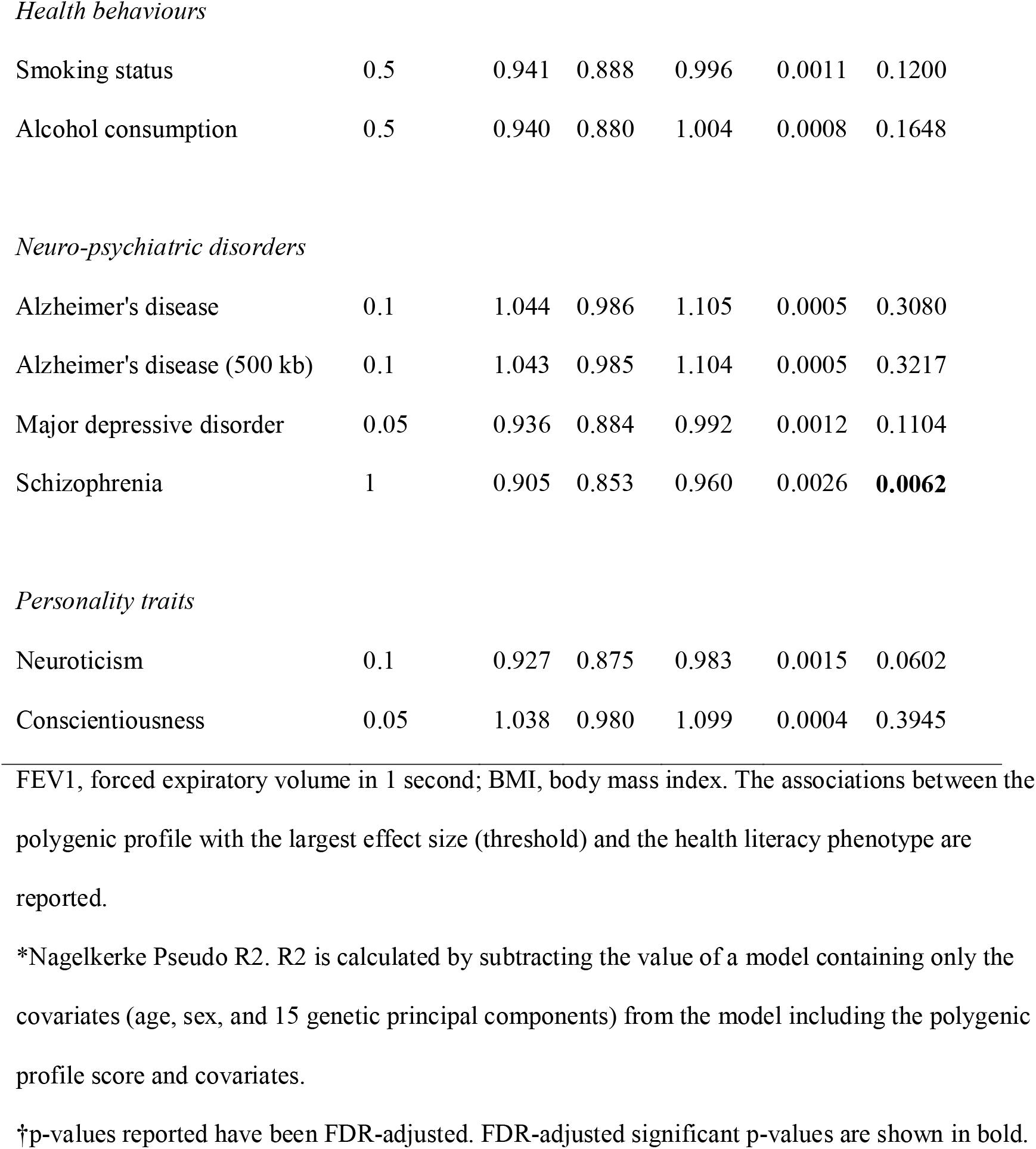
Association between polygenic profiles of cognitive, socioeconomic, health and personality traits with having adequate health literacy, controlling for age, sex, and 15 genetic principal components

|  | Threshold | OR | 95% CI |  | R2* | p-value† |
| --- | --- | --- | --- | --- | --- | --- |
|  |  |  | Lower | Upper |  |  |
| *Cognitive traits and proxies* |  |  |  |  |  |  |
| General cognitive ability | 1 | 1.339 | 1.261 | 1.422 | 0.0219 | **3.67×10<sup>-15</sup>** |
| Verbal-numerical reasoning | 0.5 | 1.304 | 1.228 | 1.385 | 0.0179 | **3.67×10<sup>-15</sup>** |
| Reaction time | 0.5 | 0.954 | 0.902 | 1.010 | 0.0006 | 0.2504 |
| Childhood IQ | 1 | 1.061 | 1.001 | 1.123 | 0.0010 | 0.1281 |
| Years of schooling | 0.1 | 1.285 | 1.212 | 1.362 | 0.0171 | **3.67×10<sup>-15</sup>** |
| *Socioeconomic measures* |  |  |  |  |  |  |
| Social deprivation | 0.01 | 0.970 | 0.916 | 1.028 | 0.0003 | 0.4530 |
| *General health measures* |  |  |  |  |  |  |
| Self-rated health | 0.1 | 0.923 | 0.871 | 0.977 | 0.0018 | **0.0359** |
| FEV1 | 1 | 0.932 | 0.879 | 0.988 | 0.0013 | 0.0880 |
| Longevity | 0.05 | 0.970 | 0.915 | 1.029 | 0.0002 | 0.4530 |
| Gait speed | 1 | 1.003 | 0.946 | 1.064 | 0.0000 | 0.9720 |
| BMI | 0.5 | 0.942 | 0.889 | 0.998 | 0.0010 | 0.1281 |
| Waist-to-hip ratio | 0.5 | 0.968 | 0.913 | 1.025 | 0.0003 | 0.4417 |
| *Chronic diseases* |  |  |  |  |  |  |
| Type 2 diabetes | 0.05 | 1.043 | 0.984 | 1.105 | 0.0005 | 0.3217 |
| High blood pressure | 0.1 | 1.056 | 0.997 | 1.118 | 0.0008 | 0.1648 |
| *Health behaviours* |  |  |  |  |  |  |
| Smoking status | 0.5 | 0.941 | 0.888 | 0.996 | 0.0011 | 0.1200 |
| Alcohol consumption | 0.5 | 0.940 | 0.880 | 1.004 | 0.0008 | 0.1648 |
| *Neuro-psychiatric disorders* |  |  |  |  |  |  |
| Alzheimer's disease | 0.1 | 1.044 | 0.986 | 1.105 | 0.0005 | 0.3080 |
| Alzheimer's disease (500 kb) | 0.1 | 1.043 | 0.985 | 1.104 | 0.0005 | 0.3217 |
| Major depressive disorder | 0.05 | 0.936 | 0.884 | 0.992 | 0.0012 | 0.1104 |
| Schizophrenia | 1 | 0.905 | 0.853 | 0.960 | 0.0026 | **0.0062** |
| *Personality traits* |  |  |  |  |  |  |
| Neuroticism | 0.1 | 0.927 | 0.875 | 0.983 | 0.0015 | 0.0602 |
| Conscientiousness | 0.05 | 1.038 | 0.980 | 1.099 | 0.0004 | 0.3945 |

FEV1, forced expiratory volume in 1 second; BMI, body mass index. The associations between the polygenic profile with the largest effect size (threshold) and the health literacy phenotype are reported.
\*Nagelkerke Pseudo R<sup>2</sup>. R<sup>2</sup> is calculated by subtracting the value of a model containing only the covariates (age, sex, and 15 genetic principal components) from the model including the polygenic profile score and covariates.
†p-values reported have been FDR-adjusted. FDR-adjusted significant p-values are shown in bold.

Increased odds of having adequate, compared to limited, health literacy were associated with a one SD higher polygenic profile score for general cognitive ability (OR = 1.34, 95% CI 1.26-1.42), verbal-numerical reasoning (OR = 1.30, 95% CI 1.23-1.39), and years of schooling (OR = 1.29, 95% CI 1.21-1.36). Reaction time and childhood IQ polygenic scores did not predict health literacy. Decreased odds of having adequate health literacy were associated with a one SD higher polygenic profile score for poorer self-rated health (OR = 0.92, 95% CI 0.87-0.98) and schizophrenia (OR = 0. 91, 95% CI 0.85-0.96). No other polygenic scores predicted health literacy.

To examine whether each polygenic profile score improved the prediction of health literacy, the Nagelkerke pseudo R2 value for a model with only the covariates (age, sex, and 15 genetic principal components) was subtracted from the Nagelkerke pseudo R2 for the model with both covariates and the polygenic score (Table 1). Polygenic profile scores for general cognitive ability, verbal-numerical reasoning, and years of schooling accounted for 2.2%, 1.8% and 1.7%, respectively, of the variance in health literacy. The variance in health literacy accounted for by the self-reported health and schizophrenia polygenic scores was small, at 0.2% and 0.3%, respectively.

Table 2 shows the results of the multivariate logistic regression in which polygenic scores for general cognitive ability, verbal-numerical reasoning, years of schooling, self-rated health, and schizophrenia were all entered simultaneously. The ORs for all polygenic scores were attenuated in this model. Increased odds of having adequate, compared to limited, health literacy were significantly associated with the following: higher polygenic scores for general cognitive ability (OR = 1.18, 95% CI 1.06-1.32), and years of schooling (OR = 1.19, 95% CI 1.11-1.27); and lower polygenic risk for schizophrenia (OR = 0.93, 95% CI 0.88-0.99). Together, these polygenic profile scores accounted for 3.0% of the variance in health literacy. In this multivariate model, the association between the verbal-numerical reasoning polygenic profile score and health literacy was attenuated and non-significant. This is not surprising as the general cognitive ability polygenic score is derived from a meta-analysis, which includes the verbal-numerical reasoning test.^44^ The self-rated health polygenic score was also attenuated and non-significant in this model.

**Table 2.** Multivariate models of the association between polygenic profiles of cognitive and health traits with having adequate health literacy, controlling for age, sex, and 15 genetic principal components

|  | OR | 95% CI |  | p-value* |
| --- | --- | --- | --- | --- |
|  |  | Lower | Upper |  |
| General cognitive ability | 1.182 | 1.058 | 1.320 | **0.0076** |
| Verbal-numerical reasoning | 1.062 | 0.953 | 1.184 | 0.3489 |
| Years of schooling | 1.186 | 1.112 | 1.265 | **9.50×10<sup>-7</sup>** |
| Self-rated health | 0.987 | 0.930 | 1.048 | 0.6691 |
| Schizophrenia | 0.929 | 0.875 | 0.987 | **0.0287** |

ORs and 95% CIs are from a model in which all five polygenic scores are entered simultaneously, controlling for age, sex, and 15 genetic principal components.
\*p-values reported have been FDR-adjusted (after a false discovery rate correction across 5 tests).
FDR-adjusted significant p-values are shown in bold.

## Discussion

Using a sample of 5,783 middle-aged and older adults living in England, no SNPs were found to be significantly associated with health literacy; however, we report 131 suggestive SNP associations within 36 independent genomic loci. Using polygenic profile scoring, this study found that genetic variants previously associated with higher general cognitive ability, verbal-numerical reasoning, and more years of schooling were associated with having adequate health literacy, whereas genetic variants previously found to be associated with poorer self-rated health and a diagnosis schizophrenia were associated with having limited health literacy. These results suggest that the phenotypic associations frequently reported between health literacy and cognitive function, and health literacy and health might be partly due to shared genetic aetiology. In a multivariate model, in which all the significant polygenic scores were entered simultaneously, higher polygenic scores for general cognitive ability, years of schooling, and schizophrenia remained significant, suggesting these polygenic scores independently contribute to performance on a health literacy test.

A number of studies have reported phenotypic associations between performance on tests of health literacy and cognitive function.^31–34^ Due to the strength of these reported associations, some researchers^33, 34^ have proposed that health literacy and cognitive function are not separate constructs, and are instead, assessing to a substantial extent the same underlying ability. To investigate this overlap, Reeve and Basalik^34^ entered three health literacy tests and six cognitive tests into an exploratory factor analysis. No unique health literacy factor emerged, and in fact, the three health literacy tests each loaded on different factors.^34^ The authors concluded that there is very little evidence that health literacy is unique from cognitive function.^34^ The current study found that the genetic variants associated with cognitive function make significant contributions to performance on tests of health literacy, providing additional evidence that health literacy and cognitive function are intrinsically related and that they might, in part, be associated with the same underlying construct.

Some researchers have suggested that educational attainment can be used as a proxy for cognitive ability in genetic studies^57, 69^ because, a) there are large phenotypic and genetic correlations between cognitive function and educational attainment,^45^ and b) it is much easier to collect information on educational attainment than it is to administer cognitive assessments in large studies. In the current study, when all significant polygenic scores were entered simultaneously, the general cognitive ability polygenic score and the years of schooling polygenic score both had independent associations with health literacy. Thus, at least when measuring health literacy, it might not be appropriate to consider cognitive function and educational attainment polygenic scores as proxies for the same underlying ability. On the other hand, it is possible that educational attainment was indexing some aspects of cognitive function not tapped by the phenotypes that went into the cognitive GWAS, which tended to be more fluid in characterisation.

The results of the current study provide some evidence that the frequently reported associations between health literacy and health^4^ might be partly due to shared genetic influences. We found that genetic variants associated with poorer self-reported health, and having a diagnosis of schizophrenia were associated with having poorer health literacy. Many studies have reported phenotypic associations between health literacy and self-reported health status.^3–5^ There has been relatively little research investigating health literacy and schizophrenia; however, health literacy has been found to be negatively associated with other mental health outcomes including mental health status^5^ and depressive symptoms.^6^ The ELSA sample used here consisted of relatively healthy community-dwelling adults. In this sample of participants without schizophrenia, having a higher polygenic risk of schizophrenia was associated with poorer health literacy. This mimics the results seen for schizophrenia and cognitive function. Individuals with higher polygenic risk of schizophrenia tend to perform more poorly on tests of cognitive function.^45, 70^ In this study, whereas the association between polygenic risk of schizophrenia and poorer health literacy was attenuated when also controlling for cognitive polygenic scores, polygenic risk of schizophrenia remained a significant predictor of health literacy, suggesting the associations reported here are not simply because of any overlap between cognitive function and schizophrenia.

A strength of the current study is that we use GWAS summary results from a large number of cognitive and health-related traits which enabled a comprehensive investigation of the shared genetic influences between health literacy, cognitive function and health. Whereas phenotypic associations between health literacy and health-related traits such as type 2 diabetes^5^ and Alzheimer’s disease^7, 8^ have been identified, we did not find that genetic variants previously associated with these health-related traits were associated with health literacy in this study. One limitation of the current study is that the quality of the polygenic profile scores created depend on the quality of the original GWAS. Many of the GWAS are meta-analyses which introduces heterogeneity in both the genetic methods used and in measuring the phenotype. Some of the GWAS have relatively small sample sizes. It is possible that we did not find an association between some of the health and cognitive polygenic scores with health literacy because the original GWAS were underpowered to identify genetic associations with the phenotype.

Unlike recent GWAS of cognitive function,^43, 44^ which found many genetic variants associated with cognitive function, we found no SNPs significantly associated with health literacy. It is now well known that, for polygenic traits, the effect of individual genetic variants on a trait are likely to be very small and, therefore, larger sample sizes than the one used here are required to identify such associations.^42^ Identification of many genetic variants associated with cognitive function are only now possible because of the ever-increasing sample sizes. The most recent GWAS of cognitive function uses data from over 300,000 individuals.^44^ The GWAS reported here is therefore underpowered.

There are few large studies that measure health literacy. This is the first investigation of the genetic contributions to health literacy. We encourage other groups with both health literacy and genetic data to explore the genetic associations of health literacy. In an effort to increase power, future studies should look to conduct a meta-analysis of GWAS of health literacy.

One strength of this study is that health literacy was measured consistently in all participants. One limitation is that the health literacy measure used in ELSA is a brief, four-item test that has a ceiling effect. That is, 70% of participants scored full marks (4/4) on this test. Despite the brief nature of the ELSA health literacy test, and despite the ceiling effect, this measure has been found to be associated with various health outcomes, including mortality.^10^ In the current study, this health literacy test was sensitive enough to identify associations with polygenic scores for cognitive and health-related traits.

Health literacy—the skills and ability required to manage ones health—have been consistently associated with cognitive function and health. This study investigated the genetic contributions to health literacy, and tested whether genetic contributions to cognitive function and health are associated with health literacy. No SNPs had genome-wide significant associations with health literacy. Polygenic scores for cognitive function, years of schooling, self-reported health and schizophrenia were associated with performance on a brief test of health literacy. These results indicate that the phenotypic associations between health literacy and cognitive function, and health literacy and health may be partly due to shared genetic aetiology between these traits. Larger studies are needed to better understand the genetic associations between health literacy, cognitive ability and health.

## Acknowledgements

The English Longitudinal Study of Ageing is jointly run by University College London, Institute for Fiscal Studies, University of Manchester and National Centre for Social Research. Genetic analyses have been carried out by UCL Genomics and funded by the Economic and Social Research Council and the National Institute on Aging. All GWAS data has been deposited in the European Genome-phenome Archive. Data governance was provided by the METADAC data access committee, funded by ESRC, Wellcome, and MRC. (2015-2018: Grant Number MR/N01104X/1 2018-2020: Grant Number ES/S008349/1).

The present study was supported by the University of Edinburgh Centre for Cognitive Ageing and Cognitive Epidemiology, part of the cross council Lifelong Health and Wellbeing Initiative, funded by the Biotechnology and Biological Sciences Research Council (BBSRC), and Medical Research Council (MRC) (grant number MR/K026992/1). The Lothian Birth Cohort 1936 is funded by Age UK (Disconnected Mind grant). SPH is funded by the Medical Research Council (MR/S0151132). This study presents independent research part supported by the National Institute for Health Research (NIHR) Biomedical Research Centre at South London and Maudsley NHS Foundation Trust and King’s College London. The views expressed are those of the author(s) and not necessarily those of the NHS, NIHR, Department of Health or King’s College London.

