## Supplementary Information for "Genetic Contributions To Health Literacy"

### Sources of genetic results from genome-wide association studies

#### AlcGen/CHARGE+

Alcohol consumption data were obtained from the AlcGen/CHARGE+ consortium.

#### CHARGE-Aging and Longevity

Gait speed data were obtained from the CHARGE-Aging and Longevity consortium.

Longevity data have been provided by the CHARGE-Aging and Longevity consortium. Longevity was defined as reaching age 90 years or older. Genotyped participants who died between the ages of 55 and 80 years were used as the control group. There were 6,036 participants who achieved longevity and 3757 participants in the control group across participating studies in the discovery meta-analysis.

Broer L, Buchman AS, Deelen J, Evans DS, Faul JD, Lunetta KL, Sebastiani P, Smith JA, Smith AV, Tanaka T, Yu L, Arnold AM, Aspelund T, Benjamin EJ, De Jager PL, Eirkisdottir G, Evans DA, Garcia ME, Hofman A, Kaplan RC, Kardia SL, Kiel DP, Oostra BA, Orwoll ES, Parimi N, Psaty BM, Rivadeneira F, Rotter JI, Seshadri S, Singleton A, Tiemeier H, Uitterlinden AG, Zhao W, Bandinelli S, Bennett DA, Ferrucci L, Gudnason V, Harris TB, Karasik D, Launer LJ, Perls TT, Slagboom PE, Tranah GJ, Weir DR, Newman AB, van Duijn CM and Murabito JM. GWAS of Longevity in CHARGE Consortium Confirms APOE and FOXO3 Candidacy. *J Gerontol A Biol Sci Med Sci*. 2015;70:110-8.

*Acknowledgments:* The CHARGE Aging and Longevity working group analysis of the longevity phenotype was funded through the individual contributing studies. The working group thanks all study participants and study staff.

**CHARGE-Cognitive working group/COGENT**

For general cognitive function a meta-analysis excluding ELSA was conducted by the CHARGE and COGENT consortia and the summary data were made available for the present study.

**CHIC**

Childhood cognitive ability data were obtained from the CHIC consortium.

**DIAGRAM**

Type 2 diabetes data were obtained from the DIAGRAM consortium.

**GIANT**

BMI and waist-to-hip ratio data were obtained from the GIANT consortium.

**International Genomics of Alzheimer's Project (IGAP)**

Alzheimer's disease data were obtained from (IGAP)

*Material and methods:* International Genomics of Alzheimer's Project (IGAP) is a large two-stage study based upon genome-wide association studies (GWAS) on individuals of European ancestry. In stage 1, IGAP used genotyped and imputed data on 7 055 881 single nucleotide polymorphisms (SNPs) to meta-analyse four previously-published GWAS datasets consisting of 17,008 Alzheimer's disease cases and 37,154 controls (The European Alzheimer's disease Initiative – EADI the Alzheimer Disease Genetics Consortium – ADGC The Cohorts for Heart and Aging Research in Genomic Epidemiology consortium – CHARGE The Genetic and Environmental Risk in AD consortium – GERAD). In stage 2, 11,632 SNPs were genotyped and tested for association in an independent set of 8,572 Alzheimer's disease cases and 11,312 controls. Finally, a meta-analysis was performed combining results from stages 1 & 2.

*Acknowledgments:* We thank the International Genomics of Alzheimer's Project (IGAP) for providing summary results data for these analyses. The investigators within IGAP contributed to the design and implementation of IGAP and/or provided data but did not participate in analysis or writing of this report. IGAP was made possible by the generous participation of the control subjects, the patients, and their families. The i–Select chips was funded by the French National Foundation on Alzheimer's disease and related disorders. EADI was supported by the LABEX (laboratory of excellence program investment for the future) DISTALZ grant, Inserm, Institut Pasteur de Lille, Université de Lille 2 and the Lille University Hospital. GERAD was supported by the Medical Research Council (Grant n° 503480), Alzheimer's Research UK (Grant n° 503176), the Wellcome Trust (Grant n° 082604/2/07/Z) and German Federal Ministry of Education and Research (BMBF): Competence Network Dementia (CND) grant n° 01GI0102, 01GI0711, 01GI0420. CHARGE was partly supported by the NIH/NIA grant R01 AG033193 and the NIA AG081220 and AGES contract N01–AG–12100, the NHLBI grant R01 HL105756, the Icelandic Heart Association, and the Erasmus Medical Center and Erasmus University. ADGC was supported by the NIH/NIA grants: U01 AG032984, U24 AG021886, U01 AG016976, and the Alzheimer's Association grant ADGC–10–196728.

**Psychiatric Genetics Consortium**

Schizophrenia and major depressive disorder data were obtained from the Psychiatric Genetics Consortium.

**Social Science Genetic Association Consortium**

Years of schooling data were obtained from the Social Science Genetic Association Consortium.

**SpiroMeta/CHARGE-Pulmonary**

Lung function (Forced expiratory volume in 1 second) data were obtained from the SpiroMeta and CHARGE-Pulmonary consortia.

**The Genetics of Personality Consortium**

Conscientiousness data were obtained from the Genetics of Personality consortium.

**The Neale Lab**

High blood pressure data were obtained from the Neale Lab.

**Tobacco and Genetics Consortium**

Smoking status (ever smoked) data were obtained from the Tobacco and Genetics Consortium.

**Supplementary Table 1** Sources of genetic results from genome-wide association studies

| Phenotype | Source | URL | Reference | GWAS n |
|---|---|---|---|---|
| General cognitive ability | CHARGE/COGENT | | Davies et al. *Nature Communications* 2018; 9: 2098. https://doi.org/10.1038/s41467-018-04362-x | 300,486 |
| Verbal-numerical reasoning | UK Biobank | https://www.ccace.ed.ac.uk/node/335 | Davies et al. *Nature Communications* 2018; 9: 2098. https://doi.org/10.1038/s41467-018-04362-x | 168,033 |
| Reaction time | UK Biobank | https://www.ccace.ed.ac.uk/node/335 | Davies et al. *Nature Communications* 2018; 9: 2098. https://doi.org/10.1038/s41467-018-04362-x | 330,069 |
| Childhood IQ | CHIC | https://www.thessgac.org/data | Benyamin et al. *Molecular Psychiatry* 2014; 19: 253-258. https://doi.org/10.1038/mp.2012.184 | 17,989 |

| | | | | |
|---|---|---|---|---|
| Years of schooling | Social Science Genetic Association Consortium | https://www.thessgac.org/data | Okbay et al. *Nature* 2016; 533: 593. https://doi.org/10.1038/nature17671 | 293,723 |
| Social deprivation | UK Biobank | https://www.ccace.ed.ac.uk/node/335 | Hill et al. *Current Biology* 2016; 26: 3083-3089. http://dx.doi.org/10.1016/j.cub.2016.09.035 | 112,151 |
| Self-rated health | UK Biobank | https://www.ccace.ed.ac.uk/node/335 | Harris et al. *International Journal of Epidemiology* 2017; 46: 994-1009. https://doi.org/10.1093/ije/dyw219 | 111,749 |
| Forced expiratory volume in 1 second ($FEV_1$) | SpiroMeta/CHARGE -Pulmonary | https://grasp.nhlbi.nih.gov/FullResults.aspx | Soler Artigas et al. *Nature Genetics* 2011; 43: 1082. https://doi.org/10.1038/ng.941 | 48,201 |
| Longevity | CHARGE-Aging and Longevity working group | https://grasp.nhlbi.nih.gov/FullResults.aspx | Broer et al. *Journals of Gerontology Series A: Biomedical Sciences and* | 6,036 cases 3,757 controls |

| | | | | |
|---|---|---|---|---|
| | | | *Medical Sciences* 2015; 70: 110-118. https://doi.org/10.1093/gerona/glu166 | |
| Gait speed | CHARGE-Aging and Longevity working group | https://grasp.nhlbi.nih.gov/FullResults.aspx | Ben-Avraham et al. *Aging* 2017: 9: 209. https://doi.org/10.18632/aging.101151 | 31,478 |
| BMI | GIANT | https://portals.broadinstitute.org/collaboration/giant/index.php/GIANT_consortium_data_files | Locke et al. *Nature* 2015: 518: 197-206. https://doi.org/10.1038/nature14177 | 339,224 |
| Waist-to-hip ratio | GIANT | http://portals.broadinstitute.org/collaboration/giant/index.php/GIANT_consortium_data_files | Shungin et al. *Nature* 2015; 518: 187-196. https://doi.org/10.1038/nature14132 | 224,459 |
| Type 2 diabetes | DIAGRAM | http://www.diagram-consortium.org/downloads.html | Scott et al. *Diabetes* 2017; *66:* 2888-2902. https://doi.org/10.2337/db16-1253 | 26,676 cases 132,532 controls |
| High blood pressure | Neale Lab | https://docs.google.com/spreadsheets/d/1b3oGI2lUt57BcuHttWaZotQcI0- | http://www.nealelab.is/blog/2017/7/19/rapid-gwas-of-thousands-of-phenotypes-for-337000-samples-in-the-uk-biobank | 336,683 |

| | | | | |
|---|---|---|---|---|
| | | mBRPyZihz87Ms_No/edit#gid=120962814 2 | | |
| Smoking status | Tobacco and Genetics Consortium | https://www.med.unc.edu/pgc/results-and-downloads | Furberg et al. *Nature Genetics* 2010: 42: 441-447. https://doi.org/10.1038/ng.571 | 74,053 |
| Alcohol consumption | AlcGen/CHARGE + | https://grasp.nhlbi.nih.gov/FullResults.aspx | Schumann et al. *PNAS* 2016: 113: 14372-14377. https://doi.org/10.1073/pnas.1611243113 | 70,460 |
| Alzheimer's disease | International Genomics of Alzheimer's Project (IGAP) | http://web.pasteur-lille.fr/en/recherche/u744/igap/igap_download.php | Lambert et al. *Nature Genetics* 2013; 45: 1452-1458. https://doi.org/10.1038/ng.2802 | 17,008 cases 37,154 controls |
| Major depressive disorder | Psychiatric Genetics Consortium (PGC) | https://www.med.unc.edu/pgc/results-and-downloads/downloads | Wray et al. *Nature Genetics* 2018; 50: 668-681. https://doi.org/10.1038/s41588-018-0090-3 | 135,458 cases 344,901 controls |

| | | | | |
|---|---|---|---|---|
| Schizophrenia | Psychiatric Genetics Consortium (PGC) | https://www.med.unc.edu/pgc/results-and-downloads/downloads | Ripke et al. *Nature* 2014; 511: 421-427. https://doi.org/10.1038/nature13595 | 36,989 cases 113,075 controls |
| Neuroticism | UK Biobank | https://www.ccace.ed.ac.uk/node/335 | Luciano et al. *Nature Genetics* 2018; 50: 6-11. https://doi.org/10.1038/s41588-017-0013-8 | 329,821 |
| Conscientiousness | Genetics of Personality Consortium (GPC) | http://eagle-i.itmat.upenn.edu/sweet/provider?uri=http://eagle-i.itmat.upenn.edu/i/00000155-e19e-5a51-c956-e86e80000000 | de Moor et al. *Molecular Psychiatry* 2017; 17: 337-349. https://doi.org/10.1038/mp.2010.128 | 17,375 |

ELSA participants were removed and the GWAS of general cognitive ability was re-run for the purpose of this analysis (n = 286,054).

**Supplementary methods**

To investigate whether health literacy polygenic profile scores predict health literacy in an independent sample, we used data from 1,005 genotyped participants in the Lothian Birth Cohort 1936 (LBC1936) study. Polygenic profile scores were created using the health literacy GWAS results and we used these polygenic scores to predict scores on a health literacy test, general cognitive ability and years of schooling in LBC1936 participants.

*Health literacy.* The Newest Vital Sign (NVS)[1] was administered to 790 LBC1936 participants at wave 2 (mean age = 70.9, SD = 0.7). In this brief health literacy test, participants are presented with a label for a container of ice cream. The participants are then asked 6 questions about the information provided on this label. The NVS is similar in content and length to the health literacy measure used in ELSA as it is a brief test health-related reading comprehension and numeracy. The score is the number of correctly answered questions (maximum score = 6). A total of 690 participants had genotyping data and scores on the Newest Vital Sign and were used as the analytic sample in the analysis examining the association between health literacy polygenic score and Newest Vital Sign.

*Cognitive ability.* The following cognitive tests, administered at wave 1 when participants were a mean age of 69.5 (SD = 0.8), were used here: Moray House Test Number 12,[2] the Wechsler Memory Scale-III (WMS-III) Logical Memory test,[3] WMS-III Spatial Span test,[3] four-choice reaction time,[4] and verbal fluency. The Moray House Test[2] is a 45-minute test of general intelligence that includes items assessing verbal reasoning and spatial ability. Total scores were corrected for age in days at testing and converted to IQ-type scores (mean = 100, SD = 15). In the Logical Memory test,[3] participants were read a paragraph containing 25 elements. The participants had to recall the story immediately and after a delay. The score used here is the total elements remembered across both immediate and delayed recall. The Spatial Span test[3] measures non-verbal working memory. The tester touches the top of a number of blocks in a specific sequence. The participant is then to tap the blocks in the same order, or in the reverse order. The sequences become progressively longer, until the participant is no longer able to remember the sequence. The score is the number of trials in which the participant correctly taps the sequence in the same or reversed order. In the four-choice reaction time test,[4] participants were presented with a box with a screen display on it and keys underneath. A number (1, 2, 3, or 4) appears on the screen and the participant has to push the key that corresponds to the number as quickly as possible. The score is the mean time (in milliseconds) to press the correct key. In the verbal fluency test, participants are asked to name as many words as they can beginning with the letters C, F and L. Participant are given one minute per letter and the score is the total number or words named within the time limit across the three trials.

Scores on these cognitive tests were entered into a principal component analysis. The first unrotated principal component accounted for 45.4% of the total variance and scores on this first unrotated principal component were saved and used as a measure of cognitive ability. Test loadings were: Moray House Test = 0.83; Logical memory = 0.65; Spatial Span = 0.63; Four-choice reaction time = -0.66; Verbal fluency = 0.57. A total of 934 LBC1936 participants had genotyping data and cognitive ability scores and were used as the analytic sample in the analysis examining the association between health literacy polygenic score and cognitive ability.

Educational attainment: At wave 1, LBC1936 participants were asked the number of years of full-time education they had completed. A total of 934 LBC1936 participants had genotyping and educational attainment data and were used as the analytic sample in the analysis examining the association between health literacy polygenic score and educational attainment.

**Supplementary Figure 1** Quantile-quantile plots for a) health literacy GWAS, and b) health literacy gene-based analysis

a

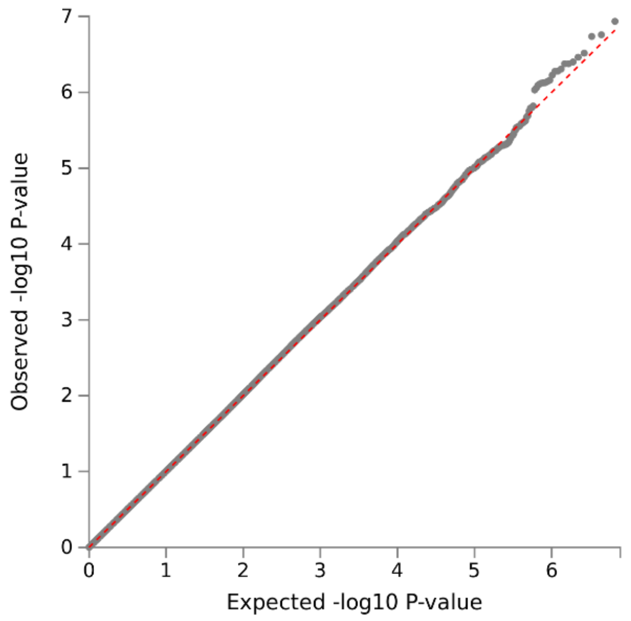


b

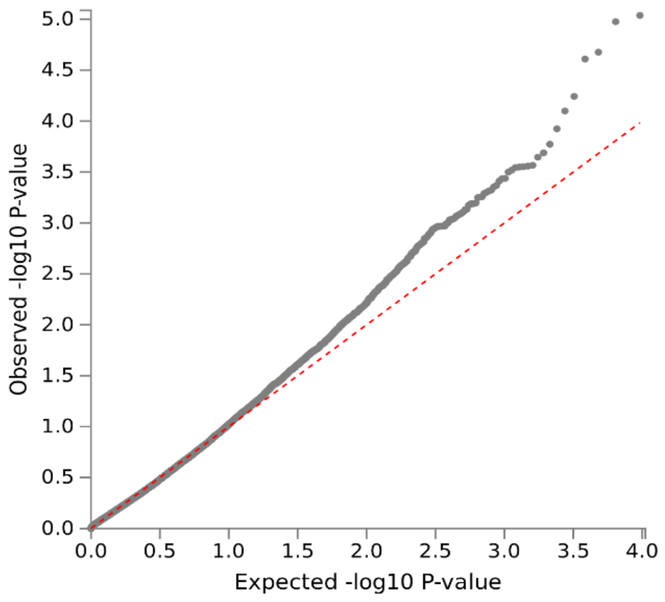

**Supplementary Table 2** Association between polygenic profile score for health literacy with the Newest Vital Sign, cognitive ability, and years of schooling, controlling for age*, sex, and 4 genetic principal components

| Trait | Threshold | β | R2 | p-value | Number of SNPs |
|---|---|---|---|---|---|
| Newest Vital Sign | 0.01 | -0.013 | -0.0013 | 0.735 | 3042 |
| | 0.05 | -0.048 | 0.0009 | 0.203 | 13091 |
| | 0.1 | -0.047 | 0.0008 | 0.217 | 24430 |
| | 0.5 | -0.020 | -0.0011 | 0.606 | 96920 |
| | 1 | -0.015 | -0.0012 | 0.693 | 152148 |
| Cognitive ability | 0.01 | 0.039 | 0.0005 | 0.219 | 3042 |
| | 0.05 | -0.004 | -0.0010 | 0.897 | 13091 |
| | 0.1 | -0.007 | -0.0001 | 0.832 | 24430 |
| | 0.5 | 0.022 | -0.0005 | 0.496 | 96920 |
| | 1 | 0.019 | -0.0007 | 0.556 | 152148 |
| Years of schooling | 0.01 | 0.060 | 0.0025 | 0.066 | 3042 |
| | 0.05 | 0.008 | -0.0010 | 0.818 | 13091 |
| | 0.1 | 0.018 | -0.0008 | 0.593 | 24430 |
| | 0.5 | 0.010 | -0.0010 | 0.766 | 96920 |
| | 1 | 0.005 | -0.0010 | 0.873 | 152148 |

R2 is calculated by subtracting the value of a model containing only the covariates (age, sex, and 4 genetic principal components) from the model including the polygenic profile score and covariates. *Age in days at wave 1 was used in the model with cognitive ability and years of schooling. Age in days at wave 2 was used in the model with Newest Vital Sign.

**Supplementary Table 3** Association between polygenic profile score for cognitive, socioeconomic, health and personality traits with having adequate health literacy, controlling for age, sex and 15 genetic principal components

| | | | | 95% CI | | | | |
|---|---|---|---|---|---|---|---|---|
| Trait category | Trait | Threshold | OR | Lower | Upper | R2* | p-value† | Number of SNPs |
| *Cognitive traits and proxies* | General cognitive ability | 0.01 | 1.283 | 1.206 | 1.365 | 0.0150 | **$2.93\times10^{-14}$** | 10644 |
| | | 0.05 | 1.302 | 1.225 | 1.384 | 0.0173 | **$3.67\times10^{-15}$** | 26498 |
| | | 0.1 | 1.332 | 1.254 | 1.416 | 0.0208 | **$2.93\times10^{-14}$** | 40550 |
| | | 0.5 | 1.339 | 1.261 | 1.422 | 0.0219 | **$3.67\times10^{-15}$** | 113154 |
| | | 1 | 1.339 | 1.261 | 1.422 | 0.0219 | **$3.67\times10^{-15}$** | 163060 |
| | Verbal-numerical reasoning | 0.01 | 1.234 | 1.157 | 1.316 | 0.0099 | **$1.03\times10^{-9}$** | 9366 |
| | | 0.05 | 1.298 | 1.219 | 1.382 | 0.0161 | **$4.92\times10^{-15}$** | 24800 |
| | | 0.1 | 1.281 | 1.204 | 1.362 | 0.0149 | **$3.30\times10^{-14}$** | 38737 |
| | | 0.5 | 1.304 | 1.228 | 1.385 | 0.0179 | **$3.67\times10^{-15}$** | 11860 |
| | | 1 | 1.302 | 1.226 | 1.383 | 0.0177 | **$3.67\times10^{-15}$** | 163279 |

| | | | | | | | |
|---|---|---|---|---|---|---|---|
| Reaction time | 0.01 | 0.999 | 0.944 | 1.057 | 0.0000 | 0.9980 | 7519 |
| | 0.05 | 0.982 | 0.928 | 1.039 | 0.0001 | 0.6334 | 21928 |
| | 0.1 | 0.976 | 0.923 | 1.034 | 0.0002 | 0.5487 | 35522 |
| | 0.5 | 0.954 | 0.902 | 1.010 | 0.0006 | 0.2504 | 109803 |
| | 1 | 0.957 | 0.904 | 1.013 | 0.0005 | 0.3048 | 163258 |
| Childhood IQ | 0.01 | 1.044 | 0.986 | 1.106 | 0.0005 | 0.3080 | 2476 |
| | 0.05 | 1.025 | 0.968 | 1.085 | 0.0002 | 0.5487 | 10086 |
| | 0.1 | 1.038 | 0.980 | 1.099 | 0.0004 | 0.3945 | 18219 |
| | 0.5 | 1.057 | 0.998 | 1.120 | 0.0009 | 0.1548 | 68420 |
| | 1 | 1.061 | 1.001 | 1.123 | 0.0010 | 0.1281 | 105819 |
| Years of schooling | 0.01 | 1.242 | 1.172 | 1.316 | 0.0129 | **$1.73\times10^{-12}$** | 8258 |
| | 0.05 | 1.262 | 1.191 | 1.337 | 0.0148 | **$4.16\times10^{-14}$** | 22772 |
| | 0.1 | 1.285 | 1.212 | 1.362 | 0.0171 | **$3.67\times10^{-15}$** | 36365 |

| | | | | | | | | |
|---|---|---|---|---|---|---|---|---|
| | | 0.5 | 1.270 | 1.198 | 1.346 | 0.0155 | **$1.36\times10^{-14}$** | 109535 |
| | | 1 | 1.265 | 1.194 | 1.341 | 0.0151 | **$2.93\times10^{-14}$** | 161726 |
| *Socioeconomic measures* | Social deprivation | 0.01 | 0.970 | 0.916 | 1.028 | 0.0003 | 0.4530 | 3910 |
| | | 0.05 | 0.980 | 0.926 | 1.038 | 0.0001 | 0.6143 | 15885 |
| | | 0.1 | 1.003 | 0.947 | 1.062 | 0.0000 | 0.9720 | 28748 |
| | | 0.5 | 1.000 | 0.945 | 1.059 | 0.0000 | 0.9980 | 106152 |
| | | 1 | 1.000 | 0.945 | 1.059 | 0.0000 | 0.9980 | 163239 |
| *General health status* | Self-rated health | 0.01 | 0.947 | 0.895 | 1.003 | 0.0008 | 0.1626 | 5052 |
| | | 0.05 | 0.932 | 0.880 | 0.986 | 0.0014 | 0.0760 | 17812 |
| | | 0.1 | 0.923 | 0.871 | 0.977 | 0.0018 | **0.0359** | 31018 |
| | | 0.5 | 0.940 | 0.888 | 0.996 | 0.0011 | 0.1200 | 107379 |
| | | 1 | 0.938 | 0.886 | 0.993 | 0.0011 | 0.1104 | 163268 |
| | FEV1 | 0.01 | 0.979 | 0.919 | 1.042 | 0.0001 | 0.6143 | 3185 |

| | | | | | | | |
|---|---|---|---|---|---|---|---|
| | 0.05 | 0.979 | 0.923 | 1.039 | 0.0001 | 0.6143 | 13275 |
| | 0.1 | 0.941 | 0.887 | 0.998 | 0.0010 | 0.1281 | 24885 |
| | 0.5 | 0.938 | 0.885 | 0.995 | 0.0011 | 0.1200 | 101307 |
| | 1 | 0.932 | 0.879 | 0.988 | 0.0013 | 0.0880 | 160612 |
| Longevity | 0.01 | 0.976 | 0.921 | 1.034 | 0.0002 | 0.5487 | 3349 |
| | 0.05 | 0.970 | 0.915 | 1.029 | 0.0002 | 0.4530 | 14278 |
| | 0.1 | 0.974 | 0.918 | 1.033 | 0.0002 | 0.5211 | 26363 |
| | 0.5 | 0.990 | 0.932 | 1.051 | 0.0000 | 0.8353 | 102168 |
| | 1 | 0.990 | 0.932 | 1.051 | 0.0000 | 0.8353 | 158944 |
| Gait | 0.01 | 1.002 | 0.945 | 1.063 | 0.0000 | 0.9848 | 3451 |
| | 0.05 | 0.991 | 0.935 | 1.051 | 0.0000 | 0.8630 | 14469 |
| | 0.1 | 0.998 | 0.941 | 1.058 | 0.0000 | 0.9858 | 26664 |
| | 0.5 | 1.000 | 0.943 | 1.060 | 0.0000 | 0.9980 | 101168 |

| | | | | | | | | |
|---|---|---|---|---|---|---|---|---|
| | | 1 | 1.003 | 0.946 | 1.064 | 0.0000 | 0.9720 | 156574 |
| | BMI | 0.01 | 0.975 | 0.921 | 1.032 | 0.0002 | 0.5211 | 4194 |
| | | 0.05 | 0.971 | 0.917 | 1.028 | 0.0002 | 0.4530 | 13316 |
| | | 0.1 | 0.965 | 0.911 | 1.022 | 0.0004 | 0.4070 | 23826 |
| | | 0.5 | 0.942 | 0.889 | 0.998 | 0.0010 | 0.1281 | 97257 |
| | | 1 | 0.948 | 0.894 | 1.004 | 0.0008 | 0.1674 | 157730 |
| | Waist-to-hip ratio | 0.01 | 0.997 | 0.941 | 1.057 | 0.0000 | 0.9720 | 3714 |
| | | 0.05 | 0.985 | 0.929 | 1.044 | 0.0001 | 0.7144 | 13918 |
| | | 0.1 | 0.968 | 0.914 | 1.026 | 0.0003 | 0.4481 | 26321 |
| | | 0.5 | 0.968 | 0.913 | 1.025 | 0.0003 | 0.4417 | 100048 |
| | | 1 | 0.971 | 0.916 | 1.029 | 0.0002 | 0.4530 | 156877 |
| *Chronic diseases* | Type 2 diabetes | 0.01 | 1.029 | 0.971 | 1.090 | 0.0002 | 0.4729 | 5265 |
| | | 0.05 | 1.043 | 0.984 | 1.105 | 0.0005 | 0.3217 | 17675 |

| | | | | | | | | |
|---|---|---|---|---|---|---|---|---|
| | | 0.1 | 1.036 | 0.977 | 1.097 | 0.0003 | 0.4263 | 31755 |
| | | 0.5 | 1.032 | 0.973 | 1.094 | 0.0003 | 0.4530 | 108370 |
| | | 1 | 1.031 | 0.973 | 1.093 | 0.0003 | 0.4530 | 163097 |
| | High blood pressure | 0.01 | 1.022 | 0.965 | 1.083 | 0.0001 | 0.5979 | 9881 |
| | | 0.05 | 1.038 | 0.980 | 1.099 | 0.0004 | 0.3945 | 25216 |
| | | 0.1 | 1.056 | 0.997 | 1.118 | 0.0008 | 0.1648 | 38804 |
| | | 0.5 | 1.033 | 0.976 | 1.095 | 0.0003 | 0.4417 | 111002 |
| | | 1 | 1.035 | 0.977 | 1.096 | 0.0003 | 0.4263 | 162052 |
| *Health behaviours* | Smoking status | 0.01 | 0.994 | 0.940 | 1.053 | 0.0000 | 0.9496 | 4065 |
| | | 0.05 | 0.997 | 0.942 | 1.055 | 0.0000 | 0.9720 | 16056 |
| | | 0.1 | 0.961 | 0.908 | 1.017 | 0.0004 | 0.3463 | 28662 |
| | | 0.5 | 0.941 | 0.888 | 0.996 | 0.0011 | 0.1200 | 102996 |
| | | 1 | 0.944 | 0.891 | 0.999 | 0.0010 | 0.1281 | 155523 |

| | | | | | | | | |
|---|---|---|---|---|---|---|---|---|
| | Alcohol consumption | 0.01 | 0.996 | 0.938 | 1.058 | 0.0000 | 0.9720 | 3806 |
| | | 0.05 | 0.979 | 0.919 | 1.042 | 0.0001 | 0.6148 | 15198 |
| | | 0.1 | 0.955 | 0.895 | 1.017 | 0.0005 | 0.3217 | 27446 |
| | | 0.5 | 0.940 | 0.880 | 1.004 | 0.0008 | 0.1648 | 101931 |
| | | 1 | 0.944 | 0.884 | 1.008 | 0.0007 | 0.2071 | 156151 |
| *Neuro-psychiatric disorders* | Alzheimer's disease | 0.01 | 1.036 | 0.979 | 1.097 | 0.0004 | 0.4046 | 4232 |
| | | 0.05 | 1.034 | 0.977 | 1.095 | 0.0003 | 0.4263 | 16247 |
| | | 0.1 | 1.044 | 0.986 | 1.105 | 0.0005 | 0.3080 | 29079 |
| | | 0.5 | 1.022 | 0.965 | 1.082 | 0.0001 | 0.5978 | 104309 |
| | | 1 | 1.016 | 0.960 | 1.076 | 0.0001 | 0.6947 | 157411 |
| | Alzheimer's disease (500 kb) | 0.01 | 1.034 | 0.977 | 1.094 | 0.0003 | 0.4263 | 4189 |
| | | 0.05 | 1.032 | 0.975 | 1.093 | 0.0003 | 0.4466 | 16192 |
| | | 0.1 | 1.043 | 0.985 | 1.104 | 0.0005 | 0.3217 | 29019 |

| | | | | | | | | |
|---|---|---|---|---|---|---|---|---|
| | | 0.5 | 1.020 | 0.964 | 1.080 | 0.0001 | 0.6143 | 104237 |
| | | 1 | 1.015 | 0.958 | 1.075 | 0.0001 | 0.7209 | 157330 |
| | Major depressive disorder | 0.01 | 1.000 | 0.938 | 1.056 | 0.0000 | 0.9711 | 5318 |
| | | 0.05 | 0.936 | 0.884 | 0.992 | 0.0012 | 0.1104 | 18589 |
| | | 0.1 | 0.940 | 0.887 | 0.996 | 0.0010 | 0.1210 | 31770 |
| | | 0.5 | 0.942 | 0.889 | 0.998 | 0.0010 | 0.1281 | 108428 |
| | | 1 | 0.937 | 0.885 | 0.993 | 0.0012 | 0.1104 | 162989 |
| | Schizophrenia | 0.01 | 0.939 | 0.882 | 1.000 | 0.0009 | 0.1356 | 9898 |
| | | 0.05 | 0.920 | 0.866 | 0.977 | 0.0018 | **0.0382** | 25557 |
| | | 0.1 | 0.922 | 0.868 | 0.979 | 0.0017 | **0.0421** | 39658 |
| | | 0.5 | 0.907 | 0.854 | 0.962 | 0.0025 | **0.0076** | 112505 |
| | | 1 | 0.905 | 0.853 | 0.960 | 0.0026 | **0.0062** | 162771 |
| *Personality traits* | Neuroticism | 0.01 | 0.939 | 0.885 | 0.996 | 0.0010 | 0.1210 | 9325 |

| | | | | | | | |
|---|---|---|---|---|---|---|---|
| | 0.05 | 0.934 | 0.881 | 0.991 | 0.0012 | 0.1030 | 24850 |
| | 0.1 | 0.927 | 0.875 | 0.983 | 0.0015 | 0.0602 | 38747 |
| | 0.5 | 0.933 | 0.881 | 0.989 | 0.0013 | 0.0921 | 111512 |
| | 1 | 0.939 | 0.886 | 0.995 | 0.0011 | 0.1200 | 163103 |
| Conscientiousness | 0.01 | 1.031 | 0.974 | 1.091 | 0.0003 | 0.4530 | 3097 |
| | 0.05 | 1.038 | 0.980 | 1.099 | 0.0004 | 0.3945 | 13315 |
| | 0.1 | 1.038 | 0.980 | 1.099 | 0.0004 | 0.3945 | 24885 |
| | 0.5 | 1.032 | 0.974 | 1.093 | 0.0003 | 0.4530 | 95148 |
| | 1 | 1.027 | 0.969 | 1.088 | 0.0002 | 0.5211 | 145404 |

***Note:*** FEV1, forced expiratory volume in 1 second; BMI, body mass index.

*Nagelkerke Pseudo R2. R2 is calculated by subtracting the value of a model containing only the covariates (age, sex, and 15 genetic principal components) from the model including the polygenic profile score and covariates.

†p-values reported have been FDR-adjusted. FDR-adjusted significant p-values are shown in bold.
